# Lipid nanoparticles incorporating a GalNAc ligand enable in vivo liver *ANGPTL3* editing in wild-type and somatic *LDLR* knockout non-human primates

**DOI:** 10.1101/2021.11.08.467731

**Authors:** Lisa N. Kasiewicz, Souvik Biswas, Aaron Beach, Huilan Ren, Chaitali Dutta, Anne Marie Mazzola, Ellen Rohde, Alexandra Chadwick, Christopher Cheng, Kiran Musunuru, Sekar Kathiresan, Padma Malyala, Kallanthottathil G. Rajeev, Andrew M. Bellinger

**Affiliations:** Verve Therapeutics, 500 Technology Square, Suite 901, Cambridge, MA, 02139, USA; Division of Cardiovascular Medicine, Department of Medicine, Perelman School of Medicine at the University of Pennsylvania, Philadelphia, PA, USA

## Abstract

Standard lipid nanoparticles (LNPs) deliver gene editing cargoes to hepatocytes through receptor-mediated uptake via the low-density lipoprotein receptor (LDLR). Homozygous familial hypercholesterolemia (HoFH) is a morbid genetic disease characterized by complete or near-complete LDLR deficiency, markedly elevated blood low-density lipoprotein cholesterol (LDL-C) levels, and premature atherosclerotic cardiovascular disease. In order to enable in vivo liver gene editing in HoFH patients, we developed a novel LNP delivery technology that incorporates a targeting ligand—*N*-acetylgalactosamine (GalNAc)—which binds to the asialoglycoprotein receptor (ASGPR). In a cynomolgus monkey (Macaca fascicularis) non-human primate (NHP) model of HoFH created by somatic knockout of the *LDLR* gene via CRISPR-Cas9, treatment with GalNAc-LNPs formulated with an adenine base editor mRNA and a guide RNA (gRNA) targeting the *ANGPTL3* gene yielded ~60% whole-liver editing and ~94% reduction of blood ANGPTL3 protein levels, whereas standard LNPs yielded minimal editing. Moreover, in wild-type NHPs, the editing achieved by GalNAc-LNPs compared favorably to that achieved by standard LNPs, suggesting that GalNAc-LNP delivery technology may prove useful across a range of in vivo therapeutic applications targeting the liver.

---

Delivery of nucleic acid therapeutics to the liver via LNPs relies on LDLR-mediated uptake into hepatocytes.^1,2^ HoFH is a genetic disease characterized by complete or near-complete absence of LDLR and, therefore, LNP treatment necessitates the use of alternative receptor pathways besides LDLR (Fig. 1). One such pathway is ASGPR, a receptor system almost entirely expressed on the surfaces of hepatocytes.^3^ Previous delivery technologies taking advantage of this receptor system include ASGPR-specific ligand conjugated antisense oligonucleotides^4,5^, siRNAs^6^, and GalNAc-LNPs constituted with siRNA^1,7,8,9,10^.

**FIGURE 1:**
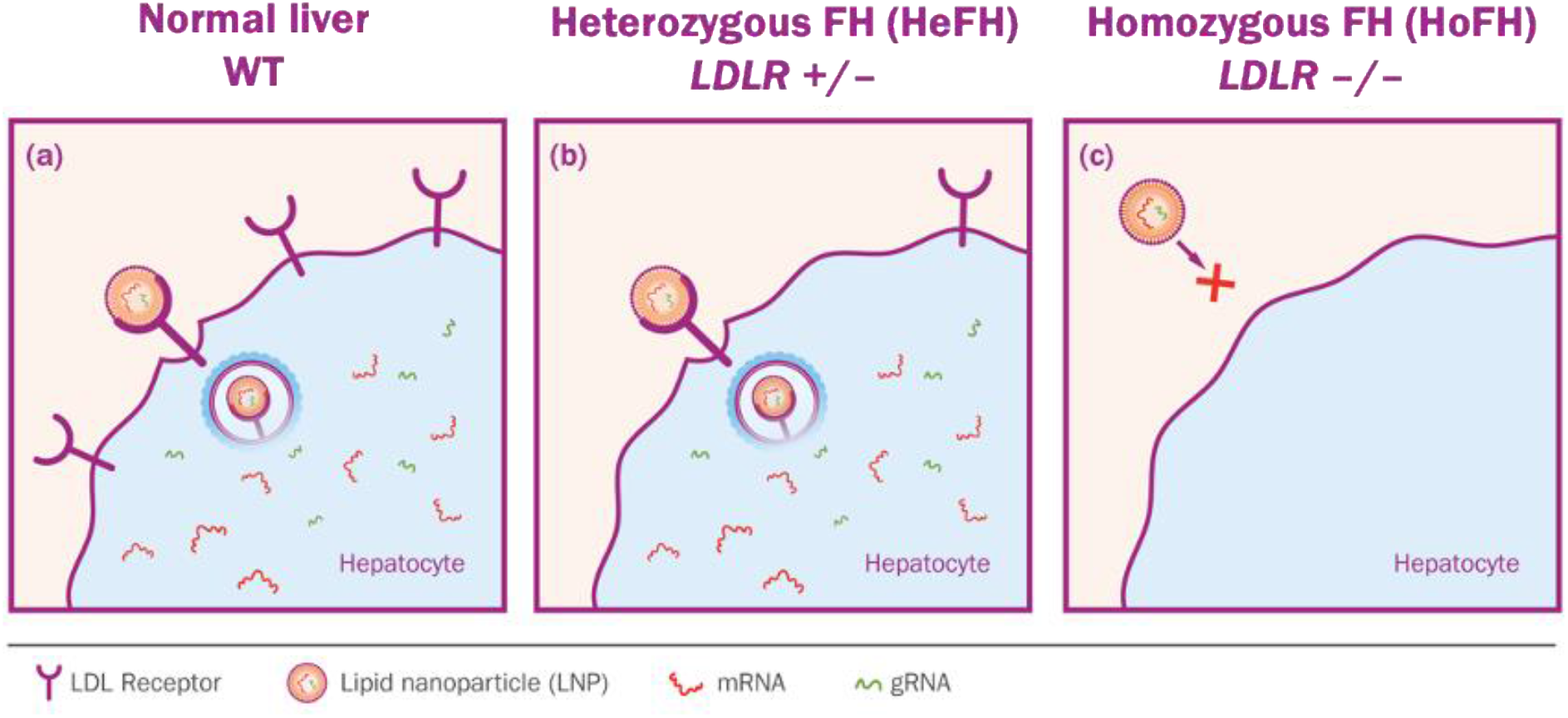
Depicts LNP uptake into hepatocytes with (a) normal LDLR expression, (b) partial deficiency as in heterozygous FH *(LDLR* +/−) and (c) complete or near-complete deficiency as may occur in homozygous FH *(LDLR* −/−). The LDLR on wild-type and *LDLR* +/− hepatocytes facilitates efficient LNP uptake into the liver, whereas the lack of LDLR on *LDLR* −/− hepatocytes stymies uptake of a standard LNP.

The work presented here expands ASGPR-mediated delivery to other nucleic acid therapeutic modalities, specifically to liver delivery of base editors^11^. We describe the development of GalNAc-LNP technology comprising a novel, potent GalNAc ligand and a scalable formulation process that together yield robust base editing in two NHP models—*LDLR* somatic knockout (KO) and wild-type (WT).

In order to enable in vivo liver base editing in the LDLR-deficient HoFH population, we designed two novel GalNAc-based ligands (Fig. 2a) and conjugated them to a series of lipid anchors (select structures shown in Fig. S1).^12^ The lysine-based trivalent ligand (Design 2) outperformed the TRIS-based ligand (Design 1) when targeting *Ldlr* −/− mice (Fig 2b, Fig. S2, Table S1). We then undertook structure-based rational design, varying the spacer length between the trivalent ligand moiety and the lipid chain, as well as incorporating three different lipid anchors (Fig. S1). We formulated GalNAc-LNPs with an adenine base editor 8.8-m (ABE8.8) mRNA and a gRNA targeting the mouse *Angptl3* gene, screened for in vivo editing activity in *Ldlr* −/− mice (Fig. S3, Table S1), and identified GalNAc-Lipid GL6 as a high performing ASGPR-targeting moiety (Fig. 2c).

**FIGURE 2:**
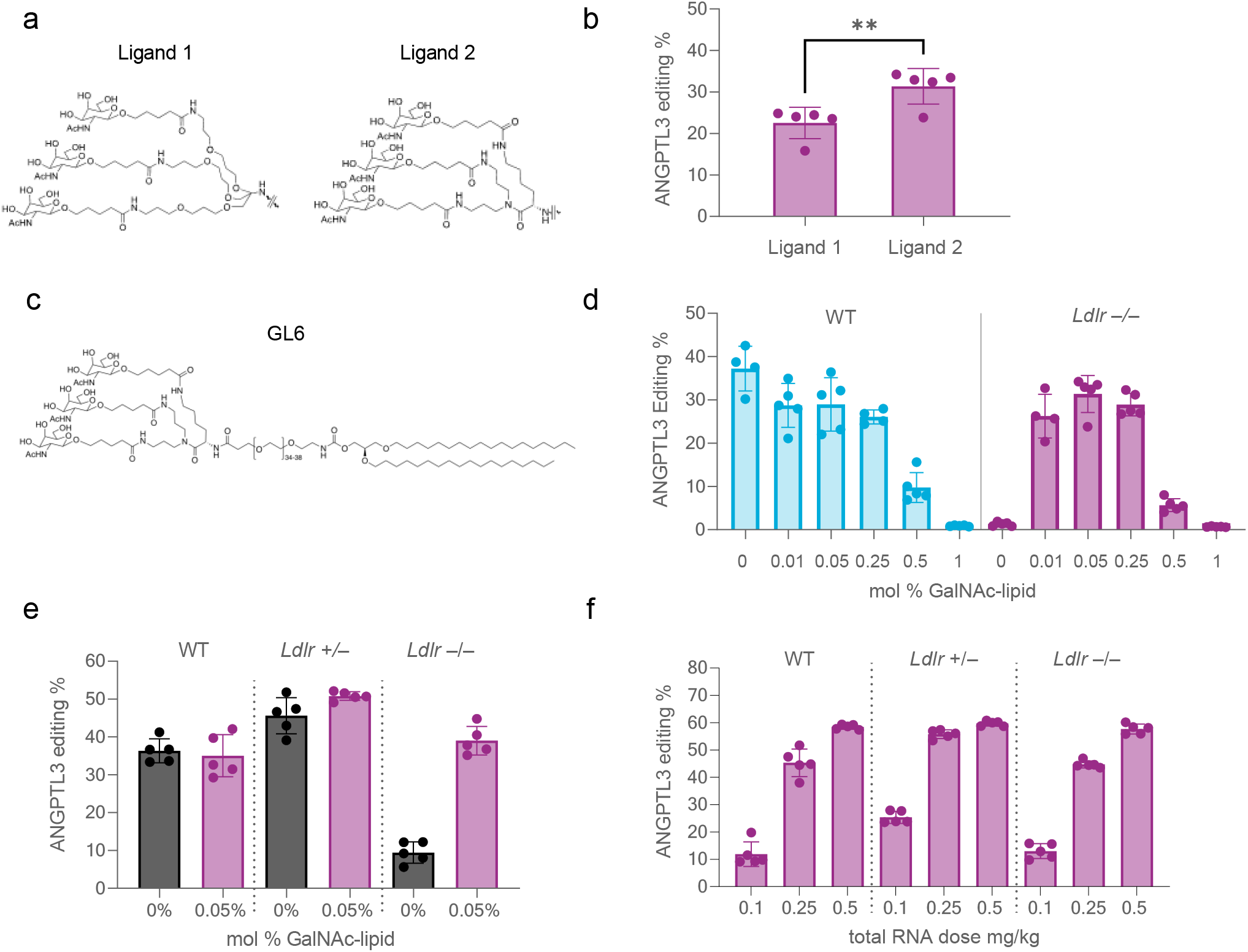
GalNAc-lipid optimization in LDLR-deficient mouse model. (a) ASGPR ligand designs 1 and 2; (b) in vivo performance in *Ldlr* −/− mice of ligand designs 1 and 2 at 0.1 mg/kg, where the PEG spacer and lipid chain were kept the same; (c) GalNAc-lipid GL6 comprising a PEG spacer and 1,2-di-*O*-octadecyl-*sn*-glyceryl lipid anchor; (d) titration of the surface density of GalNAc ligand demonstrated a low density near 0.05 mol % of GalNAc-lipid to optimally rescue liver editing in *Ldlr* −/− mice while preserving editing in WT mice at a low overall RNA dose of 0.1 mg/kg; (e) LNPs constituted with 0.05 mol % GL6 achieved rescue of *Angptl3* editing in the liver and reduction of blood ANGPTL3 protein levels in *Ldlr* −/− mice at 0.25 mg/kg; (f) demonstration of near-identical dose response of liver *Angptl3* editing using the optimized GalNAc-LNPs (constituted with 0.05 mol % GL6) in three genotypes: WT, *Ldlr* +/−, and *Lldr* −/−. Error bars represent standard deviations (s.d.), and individual data points for each animal are displayed as symbols within the bar. *n* = 5 for all panels.

Next, we developed and evaluated various options for incorporating GalNAc-lipid into the LNP during the manufacturing process. Several strategies of post-addition incorporation of GalNAc-lipid following LNP formulation resulted in a non-uniform distribution of GalNAc-lipid in the drug product based on a lectin binding assay (Fig. S4). As an alternative, we observed that mixing the GalNAc-lipid with other lipid excipients prior to LNP particle formation generated stable particles which produced similar efficacy in mice (Fig. S5) and allowed for efficient scale-up to larger batch sizes.

It has been unclear what concentration of GalNAc-lipid decorating an LNP surface would be optimal for in vivo liver delivery. LNPs targeting the *Angptl3* gene were prepared with increasing mol % of GalNAc-lipid and assessed in WT and *Ldlr* −/− mice (Fig. 2d). As little as 0.01 mol % GalNAc-lipid substantially rescued editing in *Ldlr* −/− mice without affecting Z-average size, payload entrapment, or editing performance in WT animals (Table S1). Efficacy in both mouse models remained high until about 0.3 mol % GalNAc-lipid of the total lipid content, with 0.05 mol % producing the highest editing. We confirmed the rescue of *Angptl3* editing activity of GalNAc-LNPs with 0.05 mol % GalNAc-lipid in multiple experiments across WT, *Ldlr* +/−, and *Ldlr* −/− mice (Fig. 2e, Table S1). The corresponding reductions in blood ANGPTL3 protein levels (Fig. S6) correlated well with the observed editing percentages. Dose-response comparisons in WT, *Ldlr* +/−, and *Ldlr* −/− mice demonstrated that there were near equivalent potencies in all three genotypes (Fig. 2f, Fig. S6B, Table S1) and that a 0.25 mg/kg total RNA dose with 0.05 mol % GalNAc-lipid rescued editing in *Ldlr* −/− mice and maintained editing as effectively as standard LNPs in WT and *Ldlr* +/− mice. Notably, these data indicate our ASGPR ligands may be 10-fold more potent than any GalNAc-LNP reported previously.^1,7^

Next, in order to evaluate the potency of GalNAc-LNPs in a species which might better predict human translation, we created a model of HoFH in NHPs (Fig. 3a). Two gRNAs targeting different locations on the *LDLR* gene were co-formulated with *Streptococcus pyogenes* Cas9 mRNA in a standard LNP formulation and intravenously administered to WT NHPs at a 2 mg/kg dose. With dual gRNAs targeting nearby sites in the *LDLR* gene, Cas9-mediated high-efficiency deletion of approximately 34 base pairs in the *LDLR* gene in the liver (Fig. 3b). This editing effectively knocked out liver LDLR expression (Fig. 3c) and conversely increased blood LDL-C levels about 6-fold on average (Fig. 3d), thus generating a model of HoFH in NHPs.

**FIGURE 3:**
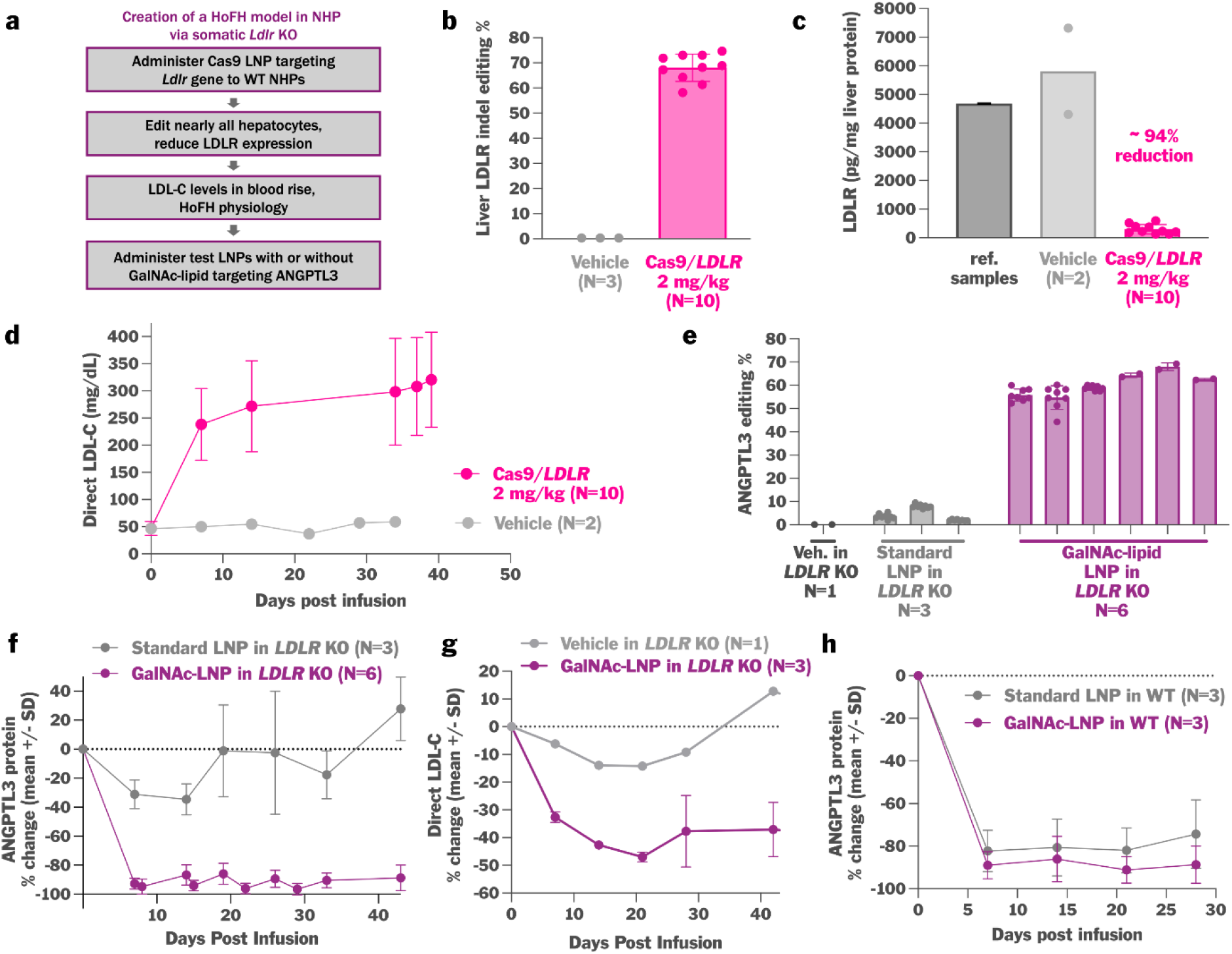
Generation of a novel somatic *LDLR* KO NHP model using CRISPR-Cas9 gene editing and successful demonstration of adenine base editing by GalNAc-LNPs targeting *ANGPTL3* in the monkey liver. (a) Schematic detailing the creation of the somatic *LDLR* KO model in NHPs. WT animals were treated with Cas9 dual gRNA LNPs targeting the *LDLR* gene, thus editing and disrupting *LDLR* in the liver. (b) A 2 mg/kg dose of Cas9 LNPs targeting *LDLR* resulted in editing of ~70% of all alleles in a liver biopsy. (c) LDLR protein levels assayed by ELISA on a second liver biopsy were markedly reduced by ~94%, and (d) blood LDL-C increased from ~50 mg/dL to ~300 mg/dL. LNPs without GalNAc-lipid were not effective in *LDLR* KO NHPs, yielding (e) <10% *ANGPTL3* editing and (f) minimal or no ANGPTL3 protein reductions at a 2 mg/kg dose. GalNAc-LNPs at a 2 mg/kg total RNA dose and with 0.05 mol % GL6 resulted in rescue of (e) high-efficiency liver *ANGPTL3* editing and (f) durable >90% reductions in blood ANGPTL3 protein. In panel (e), each column represents an NHP, and editing results reflect liver biopsy (1-2) or necropsy (8) samples. In the HoFH NHP model with markedly elevated baseline LDL-C levels of ~300 mg/dL, editing of *ANGPTL3* with the GalNAc-LNPs (g) lowered LDL-C (~35%, or ~100 mg/dL in absolute terms). (h) Comparison of blood ANGPTL3 reductions in WT NHPs dosed with either standard LNPs or GalNAc-LNPs at 2 mg/kg. Error bars represents s.d., and individual data points for each animal are displayed in panels (b), (c), and (e).

We next tested both standard LNPs and GalNAc-LNPs for the ability to deliver gene editing cargo in this HoFH NHP model. When *LDLR* KO NHPs were treated with standard LNPs (not formulated with GalNAc-lipid) (Table S2) at a dose of 2 mg/kg, minimal editing occurred at the target site in the *ANGPTL3* gene (Fig. 3e) and, thus, there was little reduction in blood ANGPTL3 protein levels (Fig. 3f). In contrast, treatment with GalNAc-LNPs (Table S2) completely rescued therapeutic efficacy in *LDLR* KO NHPs. At a 2 mg/kg dose, we observed 60% whole-liver editing of the *ANGPTL3* gene (Fig. 3e), leading to >90% lowering of blood ANGPTL3 protein (Fig. 3f), ~55% lowering of blood triglycerides (data not shown), and ~35% lowering of blood LDL-C (Fig. 3g) that in absolute terms was ~100 mg/dL of LDL-C reduction in the HoFH NHP model.

We assessed whether GalNAc-LNPs are also effective in WT NHPs in which LDLR is present. In WT NHPs, we compared GalNAc-LNPs versus standard LNPs, with each carrying ABE8.8 mRNA and a gRNA targeting *ANGPTL3*. GalNAc-LNPs achieved editing that lowered blood ANGPTL3 protein by 90%, whereas standard LNPs achieved a 75% reduction of blood ANGPTL3 (Fig. 3h). These data suggest that GalNAc-LNPs are effective across a range of genotypes including WT animals.

In summary, we have developed LNP delivery technology incorporating novel GalNAc-based ASGPR-targeting ligands and tested that technology in a NHP model of HoFH characterized by somatic LDLR deficiency in the liver. In HoFH NHPs, administration of GalNAc-LNPs carrying a adenine base editing cargo resulted in efficient editing of the target *ANGPTL3* gene in the liver, whereas standard LNPs did not. The same GalNAc-LNPs effectively delivered a base editing cargo to the livers of WT NHPs. GalNAc-LNPs provide a potent new tool for effective in vivo delivery of nucleic acid therapies.

## Notes

### Competing Interest Statement

K.M. is an advisor to and holds equity in Verve Therapeutics and Variant Bio. A.B. is an employee of Verve Therapeutics and holds equity in Verve Therapeutics, Lyndra Therapeutics, Corner Therapeutics, and Cocoon Biotech. S.K. is an employee of Verve Therapeutics, holds equity in Verve Therapeutics and Maze Therapeutics, and has served as a consultant for Acceleron, Eli Lilly, Novartis, Merck, Novo Nordisk, Novo Ventures, Ionis, Alnylam, Aegerion, Haug Partners, Noble Insights, Leerink Partners, Bayer Healthcare, Illumina, Color Genomics, MedGenome, Quest, and Medscape. All other authors are employees of and hold equity in Verve Therapeutics. Verve Therapeutics has filed for patent protection related to various aspects of therapeutic base editing of ANGPTL3 and GalNAc-lipid nanoparticle manufacture and delivery.

