## Supplemental Information and Materials and Methods for "Lipid nanoparticles incorporating a GalNAc ligand enable in vivo liver *ANGPTL3* editing in wild-type and somatic *LDLR* knockout non-human primates"

#### Standard LNPs and GalNAc-LNPs

**Table S1**: LNPs characterization and dosing data depicted in Fig. 2 and S6, which were dosed into mice via retro-orbital injection at 10 mL/kg.

| LNP | Mouse Targets[a] | GalNAc-Lipid[b] | | | Average LNP size (nm) | PDI | RNA entrapment (%) | Dose (mg/kg) | Figure |
|---|---|---|---|---|---|---|---|---|---|
| | | Compound ID | mol % | Method of incorporation | | | | | |
| 1 | *Angptl3* | GL6 | 0.05 | c | 98.61 | 0.013 | 94.91 | 0.1 | 2b |
| 2 | *Angptl3* | GL6 | 0.05 | c | 100.4 | 0.023 | 94.27 | 0.1 | 2b |
| 3 | *Pcsk9* | GL6 | 0.5 | d | 79.3 | 0.072 | 94.7 | 0.25 | S2 |
| 4 | *Pcsk9* | GL3 | 0.5 | d | 74.1 | 0.036 | 96 | 0.25 | S2 |
| 5 | *Angptl3* | GL5 | 0.05 | c | 100.6 | 0.029 | 92.48 | 0.3 | S3 |
| 6 | *Angptl3* | GL6 | 0.05 | c | 100.4 | 0.034 | 93.59 | 0.3 | S3 |
| 7 | *Angptl3* | GL7 | 0.05 | c | 98.78 | 0.032 | 92.86 | 0.3 | S3 |
| 8 | *Angptl3* | GL9 | 0.05 | c | 100.9 | 0.002 | 92.65 | 0.3 | S3 |
| 9 | *Angptl3* | GL6 | 0 | N/A | 106.6 | 0.01 | 94.85 | 0.1 | 2d |
| 10 | *Angptl3* | GL6 | 0.01 | c | 98.35 | 0.016 | 95.01 | 0.1 | 2d |
| 11 | *Angptl3* | GL6 | 0.05 | c | 98.61 | 0.013 | 94.91 | 0.1 | 2d |
| 12 | *Angptl3* | GL6 | 0.25 | c | 95.98 | 0.010 | 95.84 | 0.1 | 2d |
| 13 | *Angptl3* | GL6 | 0.5 | c | 89.63 | 0.025 | 96.15 | 0.1 | 2d |
| 14 | *Angptl3* | GL6 | 1 | c | 98.35 | 0.016 | 95.01 | 0.1 | 2d |
| 15 | *Angptl3* | GL6 | 0 | N/A | 68.09 | 0.002 | 98.66 | 0.25 | 2e, S6(A) |
| 16 | *Angptl3* | GL6 | 0.05 | c | 66.73 | 0.018 | 98.36 | 0.25 | 2e, S6(A) |
| 17 | *Angptl3* | GL6 | 0.05 | c | 77.24 | 0.028 | 99.2 | 0.1, 0.25, 0.5 | 2f, S6(B) |
| 18 | *Angptl3* | GL6 | 0.05 | d | 120.6 | 0.013 | 94.91 | 0.1 | S5 |
| 19 | *Angptl3* | GL6 | 0.05 | c | 98.61 | 0.011 | 94.27 | 0.1 | S5 |

[a] The mouse *Angptl3* and *Pcsk9* gRNAs used in these studies were selected from Chadwick *et al.* WO2021178725.
[b] The GalNAc-Lipids GL3 and GL6 were synthesized and characterized as described in Rajeev *et al.* WO2021178725.
[c] GalNAc-Lipid is premixed with other LNP excipients prior to in-line mixing with RNA to form the GalNAc-LNPs as described in WO2021178725.
[d] GalNAc-Lipid was post-inserted into the LNP to obtain the desired GalNAc-LNP (WO2021178725).
N/A: not applicable

**<u>Table S2:</u>** LNPs characterization and dosing data for generating the data shown in Fig. 3.

| LNP | NHP Target[a] | mol % GalNAc-Lipid GL6[b] | Average LNP size (nm) | PDI | RNA entrapment (%) | Dose (mg/kg) | Figure |
|---|---|---|---|---|---|---|---|
| 1 | *ANGPTL3* | 0 | 74.97 | 0.057 | 92.6 | 2 | 3e, 3f |
| 2 | *ANGPTL3* | 0 | 72.73 | 0.032 | 94.3 | 2 | 3h |
| 3 | *ANGPTL3* | 0.05 | 72.93 | 0.083 | 93.6 | 2 | 3e, 3f |
| 4 | *ANGPTL3* | 0.05 | 71.97 | 0.042 | 93.3 | 2 | 3e, 3g, 3h |

[a] The NHP *ANGPTL3* gRNA used in these studies was selected from Chadwick *et al.* WO2021178725.
[b] GalNAc-Lipid is premixed with other LNP excipients prior to in-line mixing with RNA to form the desired GalNAc-LNPs (WO2021178725).

| | |
|---|---|
| 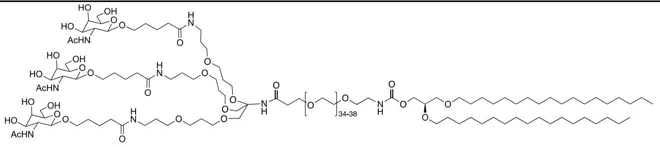 GL3 | 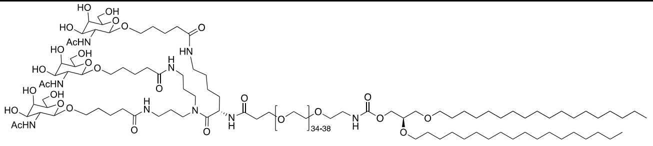 GL6 |
| 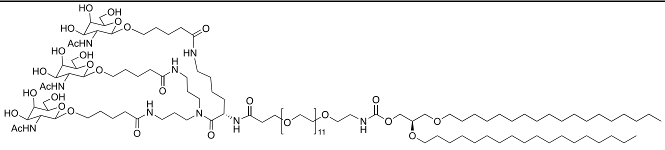 GL5 | 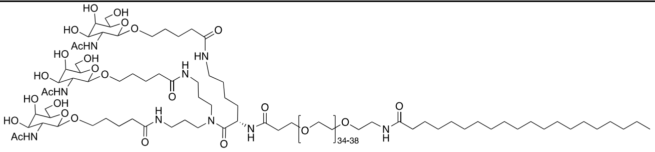 GL9 |
| 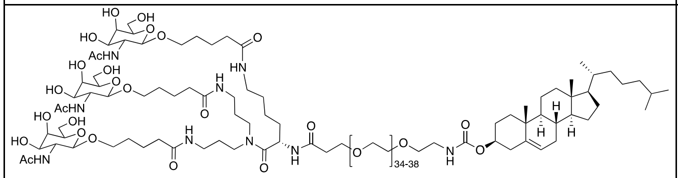 GL7 | |

**<u>Figure S1:</u>** GalNAc-based lipid ligand designs

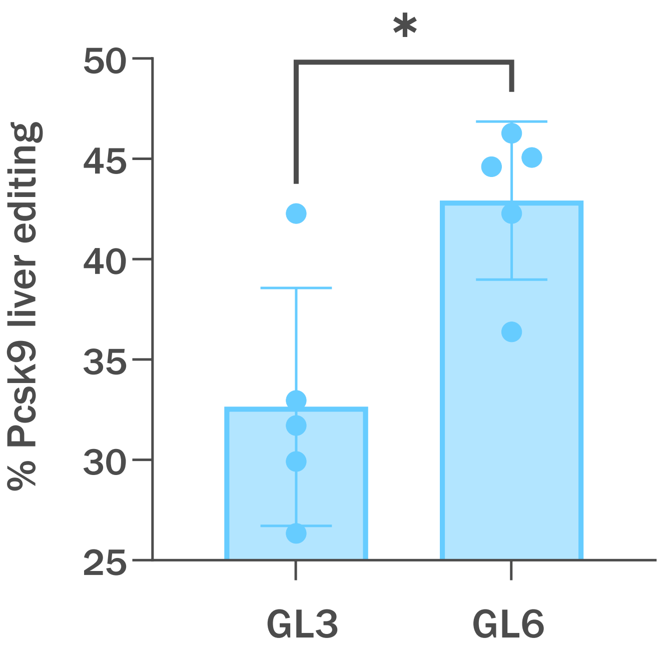


**Figure S2:** GalNAc-LNPs constituted with 0.5 mol % GL3 (Table S1, entry 4) or GL6 (Table S1, entry 3) were prepared via the post-addition method and were administered to *Ldlr* –/– mice via injection into the retro-orbital sinus. GL6 based GalNAc-LNPs achieved higher *Pcsk9* liver editing.

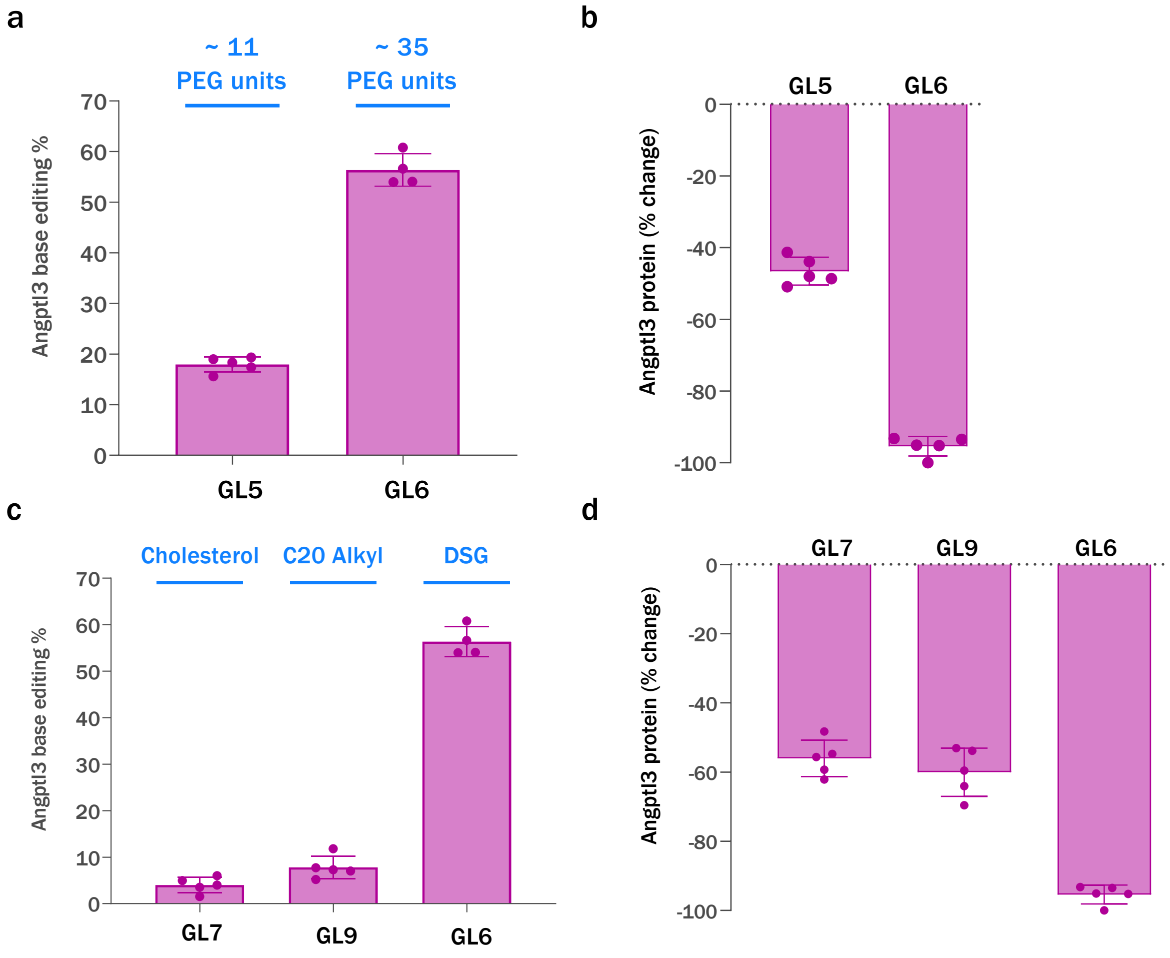


**Figure S3:** Various ligand designs were tested in *Ldlr* –/– mice at a dose of 0.3 mg/kg. GalNAc-LNPs formulated with GL6 (Table S1, entry 6) achieved a) higher editing and b) a greater reduction in ANGPTL3 protein reduction than the GalNAc-LNP with GL5 (Table S1 entry 5), the shorter PEG spacer. Modulation of the lipid tail hydrophobicity (GL7 and GL9, Table S1 entries 7 and 8) was unable to improve the c) editing efficiency and d) ANGPTL3 protein knockdown for GalNAc-LNP in *ldlr* –/– mice compared to GL6 (Table S1, entry 6).

#### Confirmation of GalNAc-Lipid Incorporation into LNP

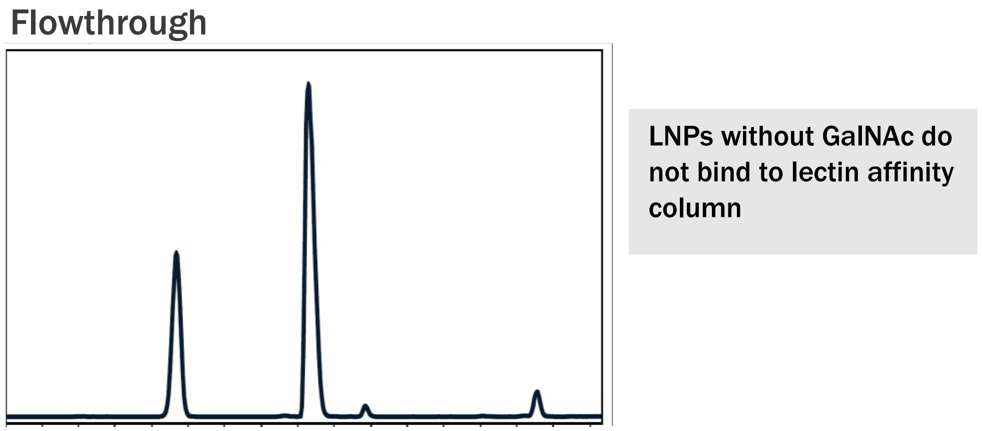


a)

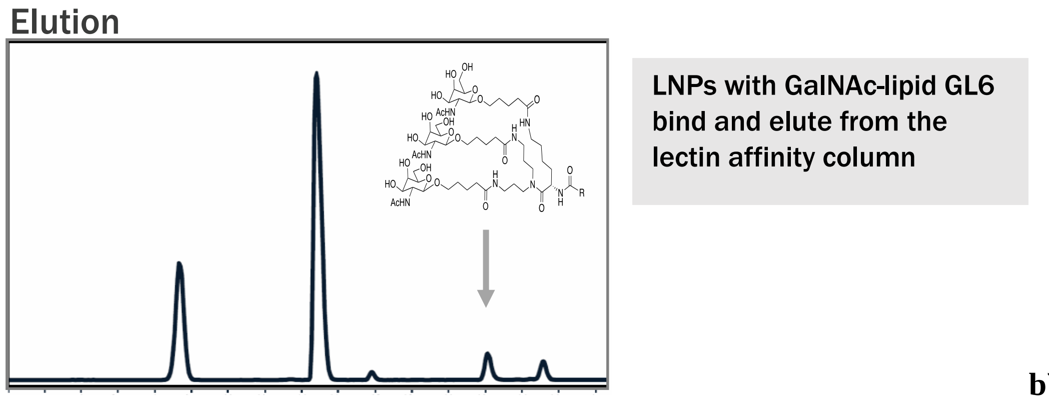


b)

**Figure S4**: We generated GalNAc-LNPs with 0.5 mol % GL6 via the post-addition method of GalNAc-lipid incorporation. These LNPs were then introduced into a lectin affinity column, with PBS flowed into the column afterwards. LNPs containing GalNAc-lipid would be expected to bind to the lectin column. Fractions of the flowthrough were then collected. If all LNPs contained GalNAc-lipid, no LNPs would be expected in this flowthrough. PBS with D-(+)-Galactose was then added to the column, and more flowthrough fractions were collected. Galactose is expected to displace GalNAc-LNPs from the column and now allow them to be eluted in the flowthrough. a) Fractions collected from the flowthrough and rinse fractions in PBS when analyzed by IP-RPLC-HPLC-ELSD showed no GalNAc-lipid but maintained the other 4 lipids in their expected molar composition. This indicates that a population of the post-addition manufactured LNPs do not actually contain GalNAc-lipid at all, as they were unable to bind to the lectin column and were flushed out by the PBS. b) Fractions collected from the elution with PBS containing D-(+)-Galactose showed GalNAc-lipid in addition to the other four LNP excipients. Some of the LNPs generated via the post-addition method did indeed contain GalNAc-lipid, and so were displaced as expected by the Galactose. Ultimately, this assay indicates that the post-addition method generates non-homogenous distributions of GalNAc-lipid, with a population of the LNPs lacking GalNAc-lipid completely, as seen in Panel a).

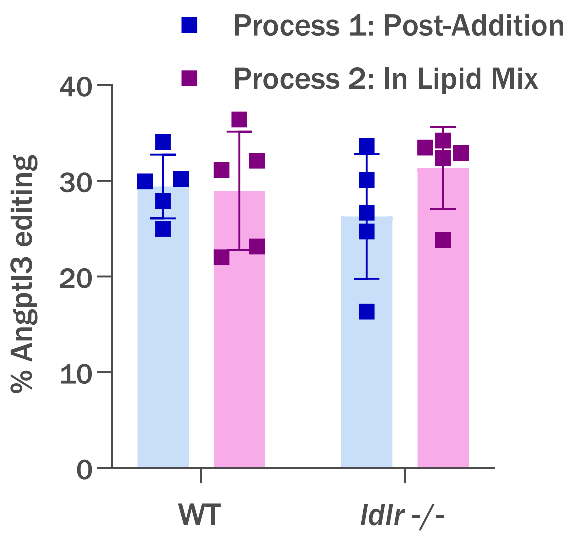


**Figure S5:** GalNAc-LNPs were manufactured with 0.05 mol % GalNAc-lipid GL6 added via post-addition or in lipid mix methods (Table S1, entries 18 and 19). GalNAc-LNPs prepared by both methods produced nearly identical editing in WT and *Ldlr* –/– mice at a 0.1 mg/kg dose.

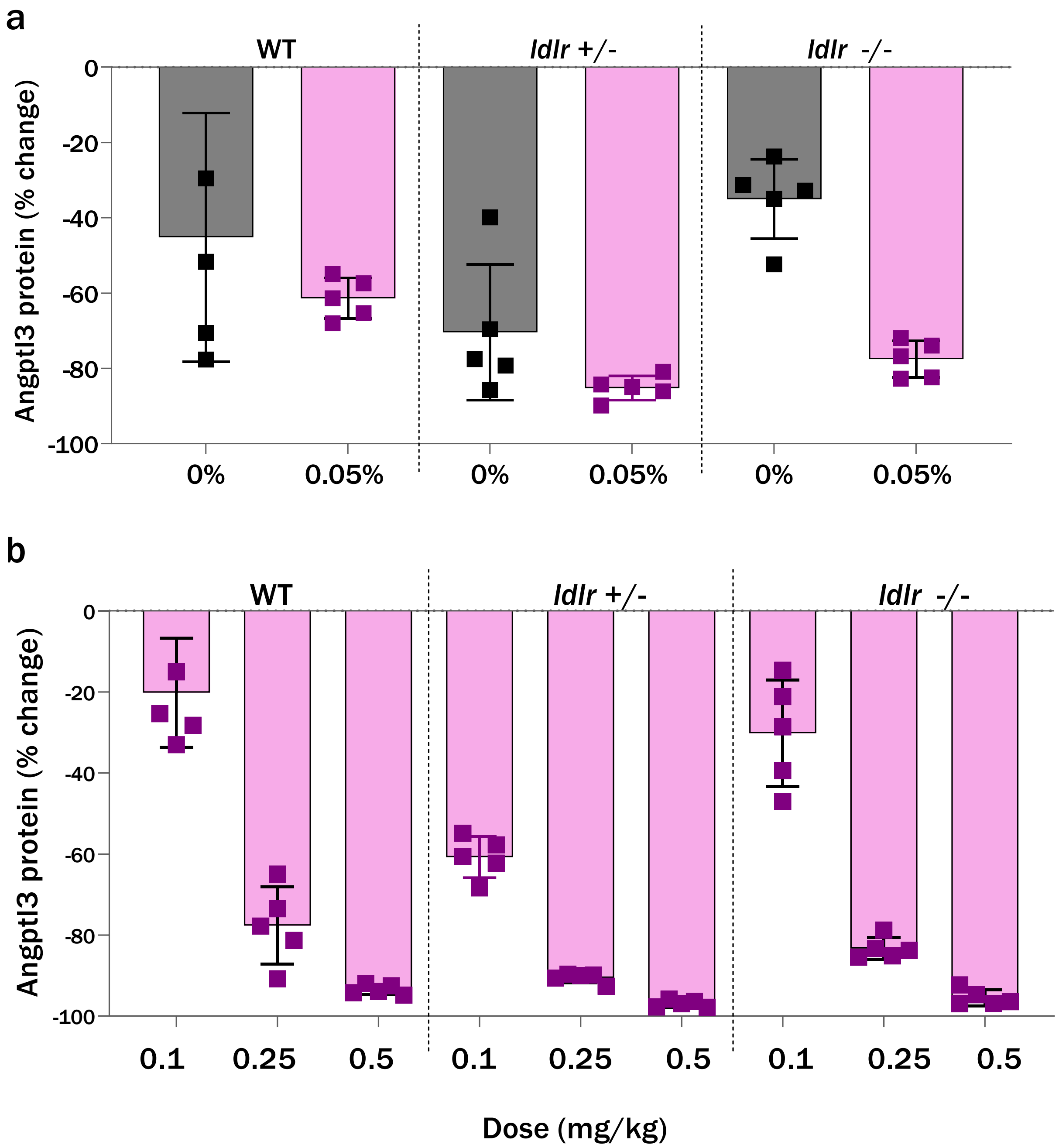


**Figure S6:** LNPs formulated with 0.05 mol% of GL6 were administered to WT, *Ldlr* +/–, and *Ldlr* -/– mice at 0.25 mg/kg in Panel A and the stated doses in Panel B. a) The inclusion of GalNAc-lipid GL6 rescued activity in L*dlr* –/– mice and maintained activity in WT and L*dlr* +/– (Table S1, entries 15 and 16). b) Dose-dependent Angptl3 protein reduction was achieved in all three animal models with the GalNAc-LNP evaluated (Table S1, entry 17).

### Materials and Methods

#### Preparation of Standard and GalNAc-LNPs

LNPs used in the studies were prepared as described in the patent publication, WO2021178725 (Tables S1 and S2). Each LNP is comprised of an ionizable amino lipid, a PEG-Lipid, cholesterol and distearoyl-sn-glycerol-3-phosphocholine (DSPC). In addition to the standard excipients each GalNAc-LNP contains a select GalNAc-lipid shown in Figure S1. The GalNAc-LNPs were prepared either by mixing the ligand conjugated lipid with the excipient prior to LNP preparation or by a post-insertion process as described in WO2021178725. The LNPs shown in Table S1 were constituted with the adenine base editor 8.8-m (ABE8.8) mRNA and a gRNA targeting the mouse *Angptl3* or *Pcsk9* gene. For the NHP studies, the LNPs were constituted with the adenine base editor 8.8-m (ABE8.8) mRNA and a gRNA targeting the monkey ANGPTL3 gene (Table S2).

#### LNP Analytics and characterization

LNP critical quality attributes – Z average size, polydispersity, total RNA concentration, encapsulation efficiency, and lipid content – were determined after particle formation. LNP size and polydispersity were measured using Dynamic Light Scattering via a Malvern Panalytical Zetasizer Ultra. Encapsulation efficiency was determined by fluorimetry using Ribogreen dye, as previously described (Geall AJ, Verma A, Otten GR, Shaw CA, Hekele A, Banerjee K. et al. Nonviral delivery of self-amplifying RNA vaccines. Proc Natl Acad Sci USA. 2012, 109:14604). For the evaluation of lipid composition of the LNPs, an ion-pairing reverse phase high performance liquid chromatography with evaporative light scattering detector (IP-RPLC-HPLC-ELSD) was used. The assay uses a standard curve to quantify the amino lipid, PEG-Lipid, cholesterol, DSPC, and GalNAc-lipid.  For the lectin affinity column-based analysis of GalNAc-LNP, the LNP was allowed to pass through a lectin affinity column. The flowthrough was collected and analyzed for unbound ligand-free LNPs. After washing the lectin column with loading buffer, the column was washed with PBS buffer containing D-(+)-galactose and the eluent were collected and analyzed in a similar fashion to evaluate the LNPs that bound to the lectin column.

**Animal studies**

Mouse studies were approved by the Institutional Animal Care and Use Committee of the Charles River Accelerator and Development Lab (CRADL) where the studies were performed. Female 8-10 weeks old C57BL/6J, *Ldlr* +/–, and *Ldlr* –/– mice from The Jackson Laboratory were used for the mouse studies, with random assignment of mice to experimental groups, and with collection and analysis of data performed in a blinded fashion. The mice were maintained on 12-h light/12-h dark cycle, with a temperature range of 65 °F to 75 °F and a humidity range of 40% to 60%. LNPs were administered to the mice via injection into the retro-orbital sinus. Five to ten days following treatment, the mice were euthanized, and liver samples were obtained on necropsy and processed with the KingFisher Flex Purification System according to the manufacturer's instructions to isolate genomic DNA.

NHP studies were approved by the Institutional Animal Care and Use Committees of Altasciences. Two similarly designed NHP studies were performed to confirm and extend the results, both used cynomolgus monkeys (*Macaca fascicularis*) of Cambodian origin. The animals were 2-3 years of age and 2-3 kilograms in weight at the time of study initiation. All animals were genotyped at the *ANGPTL3* editing site to ensure that any animals receiving ABE8.8/ANGPTL3 LNPs were homozygous for the protospacer DNA sequences matching the gRNA sequence; otherwise, animals were randomly assigned to various experimental groups. The animals were premedicated with 1 mg/kg dexamethasone, 0.5 mg/kg famotidine, and 5 mg/kg diphenhydramine on the day prior to LNP administration and then 30-60 minutes prior to LNP administration. The LNPs were administered via intravenous infusion into a peripheral vein over the course of 1 hour. Control animals received phosphate-buffered saline instead of LNPs under the same infusion conditions.

In both NHP experiments, WT NHPs were dosed with LNPs containing spCas9 and an LDLR guide RNA pair. Liver biopsies were taken at Day 19 to assess LDLR editing. Subsequently, at least 30 days after initial treatment, WT and newly generated LDLR somatic KO NHPs were injected with LNPs carrying ABE mRNA and ANGPTL3 guide RNA.  For blood chemistry samples, animals were fasted for at least 4 hours before collection via peripheral venepuncture.

In both NHP studies, samples were collected on the following schedule: day –10, day –7, day –5, day 1 (6 hours after LNP infusion), day 2, day 3, day 5, day 8, and day 15. Blood samples were analysed by the study site for LDL cholesterol, HDL cholesterol, total cholesterol, triglycerides, AST, and ALT. A portion of each blood sample was used for ANGPTL3 protein measurement. In both NHP studies, each animal underwent a liver biopsy via laparotomy on day 15 after administration of the first LNP. In one NHP study, each animal underwent euthanasia and necropsy on day 75. On necropsy, liver samples were collected by protocol. Two samples each were collected from the left, middle, right, and caudate lobes, for a total of eight samples per liver. Organ samples were processed with the KingFisher Flex Purification System (Thermo Fisher) according to the manufacturer's instructions to isolate genomic DNA.

**NGS editing and ANGPTL3 ELISA**

The DNA base editing was assessed using PCR primers specific to the targeted genomic site by following the procedure described by Musunuru et al. (*Nature*, **2021**, 593:429-434). Briefly, PCR reactions used Accuprime High Fidelity DNA Polymerase (Thermo Fisher, #12346-094) with primers specific to the target *Angptl3* genomic site with 5' Nextera adaptor sequences, followed by purification of the PCR amplicons with the Sequalprep Normalization Plate kit (Thermo Fisher, #A1051001). A second round of PCR with the Nextera XT Index Kit V2 Set A (Illumina, #15052163) and/or Nextera XT Index Kit V2 SetD (Illumina, #15052166), followed by purification of the PCR amplicons with the Sequalprep Normalization Plate kit, generated barcoded libraries, which were pooled and quantified using a Qubit 3.0 Fluorometer. After denaturation, dilution to 8 pM, and supplementation with 15% Phix (Illumina, #15017666), the pooled libraries underwent paired-end sequencing on an Illumina Miseq System.

The ANGPTL3 plasma protein levels were performed using murine or human, as appropriate, ANGPTL3-specific ELISA assay developed in our laboratory (Musunuru *et al*. *Nature*, **2021**, 593:429-434 and Chadwick *et al*. WO2021178725).
